# Pectin is a critical contributor to the mechanical properties of *Chara* cell walls

**DOI:** 10.64898/2026.09.16.751926

**Authors:** Zhe He, Junsoo Kim, Zheqi Chen, Ian D. Svetkey, Zhigang Suo, Fulton Rockwell, N. Michele Holbrook

## Abstract

Cellulose has long been viewed as the major load-bearing component in plant primary cell walls. Recent studies, however, suggest that pectin plays a significant role in cell wall mechanics despite a kPa-scaled modulus. Here we quantify the mechanical properties of centimeter-sized internodal cell walls of *Chara corallina* following treatments to remove calcium crosslinks or digest pectin. We also characterize the flow of water and gas through the cell wall as a function of pressure. We found that removal of calcium crosslinks did not affect wall strength and stress-relaxation behavior, yet it decreased the elastic modulus and increased permeability to water. In contrast, pectin removal had a dramatic impact on all measurements. Samples from which pectin had been removed exhibited lower strength and stiffness and had greater stress relaxation under constant strain. Pectin removal increased the permeability of the cell wall to water above that observed when calcium was removed and reduced the pressure at which gas “tunnels” through the cell wall ∼ 2-fold. Our findings support the idea that pectin locally restricts the movement of cellulose microfibrils, with implications for understanding functional properties of primary cell walls in vascular plants and developing high-performance double-network materials.

## I. INTRODUCTION

Primary cell walls are integral to the functioning of plants: they provide the structural integrity of the plant body, sustain the turgor pressure of living cells, and serve as a physical barrier against pathogens [1-3]. The primary cell wall is a composite material composed of several major classes of polymers, with cellulose accounting for approximately 20-40% of the dry mass, pectin for 30-50%, and hemicellulose for 20–30% [4]. Cellulose in plant cell walls is a fibrous network consisting of layers of stiff microfibrils, often described as a “multilayer nanostructure” [5]. Cellulose microfibrils are composed of 18 unbranched β-1,4-glucan chains [6], and individual microfibrils exhibit a Young’s modulus on the order of gigapascals [5]. Pectin, on the other hand, is an anionic branched polymer that can form crosslinks via calcium ions, which allows gelation to occur, yielding a hydrogel with elastic modulus almost ten times higher than the viscous modulus [7]. Hemicellulose, another branched polymer, is believed to function as a “tether” that links cellulose microfibrils together [8], although a few studies pose challenges to this view [9].

Historically, cellulose has been thought to be the major load bearing component in primary cell walls based on experiments and simulations under uniaxial loading conditions [10-12]. Pectin has a kilopascal shear modulus [13] and is thought to be more of a “filler” that largely affects porosity rather than mechanics [14-16]. Recent studies, however, suggest that pectin makes a significant contribution to the mechanics of cell walls [17-20]. Pectin is more abundant near cellulose microfibrils than hemicellulose [9], suggesting that pectin’s contribution to the mechanical properties of cell walls arises from the ways in which the hydrogel interacts with the much stiffer cellulose microfibrils.

In this study, we use uniaxial and biaxial measurements to uncover the ways in which pectin affects the mechanical properties and functioning of plant cell walls. We focus on *Chara corallina*, a freshwater green alga in the sister group (Charales) to the land plants [21], whose cell walls have been used extensively as a model system for studies of plant cell wall development. *Chara’s* internodal cells are typically 1 to 10 centimeters long and 0.6-0.8 millimeters in diameter [21,22], have minimal heterogeneity and thus are amenable to various mechanical tests. We hypothesize that pectin stabilizes the spacing of cellulose microfibrils against MPa scale stresses in the cell wall of *Chara*.

## II. METHODS

A dense colony of *Chara carollina* was grown in a large (∼0.7 × 0.5 × 0.3 m) indoor tank filled with spring water and supplied with supplemental lighting. Long and mature internodes [23] ranging from 6-9 cm were gently removed using a razor blade, with at least one node intact. If the internode contained two ends, one end of the cell was cut near the node in the transverse direction, and the cytoplasm gently expelled. Mature *Chara* cell walls are reported to be ∼ 10 µm thick [25]. This was confirmed by embedding a transverse section of cell wall in resin (NOA 63, Norland Products) while wet, sectioning the sample transversely, and visualizing under an optical microscope (Leica DM6 FS) to capture images that were analyzed in ImageJ.

Four treatment groups were prepared: (1) calcium removal, in which the calcium crosslinks in the cell walls were removed using a chelator solution containing 10mM NaPO_4_ at pH 10, and subsequently kept in de-ionized (DI) water until measurements on the same day; (2) pectin removal, in which cells were incubated for 4 h at 37 °C in 5mM MES (2-(N-morpholino)ethanesulfonic acid) at pH 5 buffer containing 0.1% (w/v) pectolyase, and kept in DI water until measurements on the same day; (3) control, in which freshly collected cells were immediately tested; and (4) control shake, in which cells were maintained in spring water, but exposed to the same mechanical perturbations as the calcium and pectin removal treatments. Samples from all groups except for the control were placed in ∼5 mL of their corresponding fluid medium in 12 mL Falcon tubes on a shaker for ∼12 h (mechanical characterizations and water permeability measurements) and for 4 h (air permeability measurements). For calcium and pectin removal samples, DI water was used for post-treatment storage to prevent calcium ions from diffusing back into the walls.

### A. Mechanical characterizations

#### 1. Tensile test

Six cells were selected from each of the treatment groups described above. As shown in FIG. S1 of the Supplemental Material [24], the cell wall was wrapped around two wooden sticks at both ends and secured with a fast-setting adhesive (Krazy Glue). Subsequently, the two wooden sticks at the specimen ends were clamped onto a tensile machine (Instron 5966, 100 N load cell) for testing. We applied water to the sample using a pipette to maintain hydration during measurements. The stretch rate was 0.05 s^-1^. From the stress-strain curves, we extracted the strength, the stress at material failure point using an estimated cross-sectional area (circumference × wall thickness), and Young’s modulus E, the slope of the linear region of the stress-strain curve, both have a unit of MPa.

#### 2. Stress-relaxation test

Cell walls from control, calcium removal, and pectin removal groups were clamped to the Instron via wooden sticks as described above. We imposed a constant stretch and recorded stress as a function of time. We normalized the stress *s* by its initial value *s*_0_, *s*/*s*_0_, and plotted the normalized stress versus time in FIG. 1(d). The applied stretch was determined by the corresponding stretch at ∼70% of the average ultimate strength of each treatment shown in FIG 1(a). The humidity was maintained at ∼100% during measurements.

**FIG. 1.**
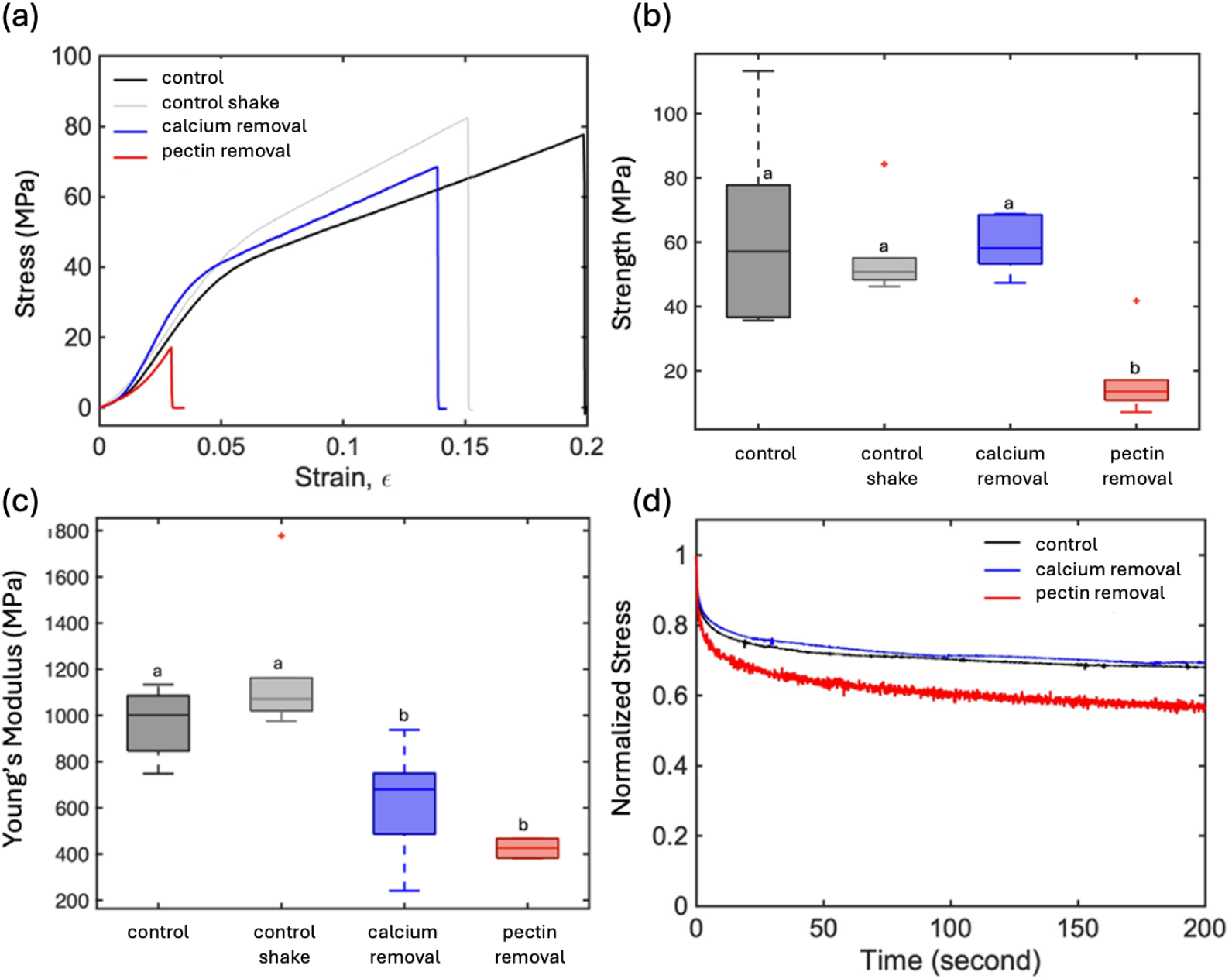
Mechanical tests on *Chara* cell walls. (a) Uniaxial tensile test (n = 6 for each treatment, one representative curve for each group are shown); (b) strength (p < 0.05, n = 6); (c) Young’s modulus (one-way ANOVA, p < 0.05, n = 6); (d) stress-relaxation tests (n=3, one representative curve for each group are shown). Error bars represent standard errors (SE), and lower-case letters show statistical difference after Tukey-adjusted pairwise comparison.

#### 3. Toughness measurement

To measure the fracture toughness of cell walls from control, calcium removal, and pectin removal treatments, a long syringe needle (0.4 x 50 mm, Henke-Ject) was inserted into the cylindrical wall of a cell cut open at both ends and used as a support in making a longitudinal cut along the length of the cell.^22^ The resulting single-layered rectangular cell wall (> 3 cm long) was carefully glued onto two pieces of PET backing layer (8567K12, McMaster, 30 μm), as shown in FIG. S2 of the Supplementary Material. The two ends of the plastic sheet were clamped onto a tensile machine (Instron 5966, 100N load cell) and then pulled at a constant rate of 1 mm s^-1^. The force first rose and then plateaued at *f*. The toughness is calculated as twice the plateau force divided by the specimen thickness *t, G*_c_ = 2*f*/*t*.

### B. Water injection experiment

We subjected the cell wall to biaxial stress by injecting water into cylindrical cells that had been emptied of their contents. Four cells from control, calcium removal, and pectin removal groups were selected, with one end sliced open and the other end capped with an intact node. The open end was attached to a glass capillary needle using Loctite 409 Instant Adhesive and Loctite SF 7452 glue accelerator, followed by curing for approximately 5 min. During the curing process, cells were kept hydrated by suspending them in water with only the glued end exposed. Afterwards, more glue was applied at the other end of the cell where the node was attached, and an accelerator was applied to reduce the curing time. After another 5 min, the capillary was filled with water subjected to incrementally increasing pressures using a compressed nitrogen tank (10 psi per step until 150 psi), and the water flow rate through the cell walls was measured using a Sensirion flow meter (SLC1430-025, Sensirion AG) shown in FIG 2(a). A complete set of water injection measurements was performed for four cells per chemical treatment.

**FIG. 2.**
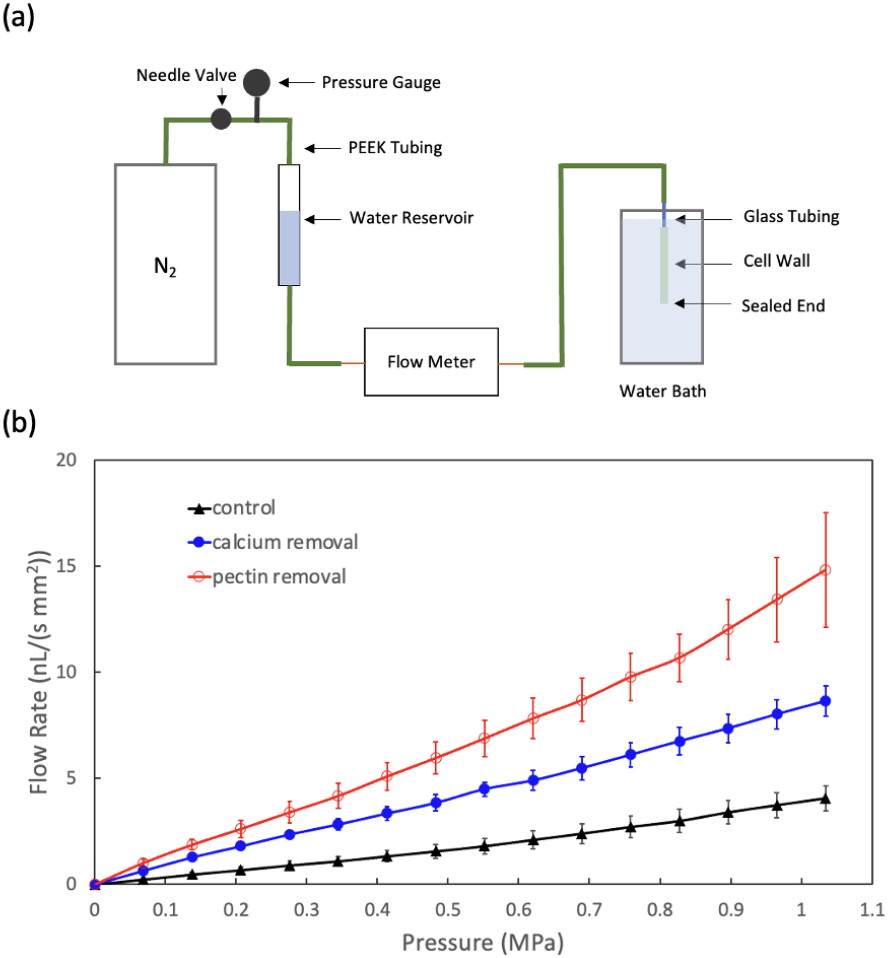
Water injection experimental setup and results. (a) Schematic of flow measurement setup: a liquid nitrogen tank is connected to a customized water reservoir via polyetheretherketone (PEEK) tubings, and water flows through the flow meter, followed by a pulled capillary glass tubing (see Materials and Methods), and out of the *Chara* cell wall. (b) Flow rate of water vs. applied pressures of *Chara* cell walls. Control (n=4), calcium removal (n=4), and pectin removal (n=4). The flow rate is normalized by the unglued sample area which is permeable to water. Error bars for the flow rate curves are standard error at a 95% confidence interval.

To ensure the integrity of the glued connections, each sample was initially filled with air and then submerged in water and pressurized to 60 psi. The absence of bubbles forming rapidly near the glued connections indicated a secure connection. Samples in which bubbles were observed were discarded. The glass capillary needles were fabricated using a microcapillary puller (PUL-1, World Precision Instruments). A calibration curve was obtained using a 50 µL syringe and a syringe pump (Harvard Apparatus, NE-300) directly connected to the Sensirion flow meter in FIG. S3 of the Supplementary Material.

### C. Gas injection experiment

We also injected air into cylindrical cells that had been emptied of their contents. Gas injection imposes biaxial stress in the cell wall while creating a gas-water interface that has a non-zero surface tension. Although gases can diffuse through the cell wall, we were interested in the pressure at which gas penetrates the cell wall, forming steady streams of bubbles on the unpressurized side. Control and pectin-removal samples were glued to glass capillaries connected to a compressed nitrogen gas tank, as described for the water permeability measurements. A 0.1 M sodium dodecyl sulfate (SDS) solution was used as an immersion bath to reduce surface tension between gas and liquid, lowering the surface energy and pressure threshold for gas to transit across the cell wall before the cell wall bursts. Samples were submerged in the surfactant solution, with the glued end above the liquid surface, for a few minutes to allow SDS to diffuse into the cell wall. Because SDS is an anionic charged surfactant, it is not expected to affect pectin mechanical properties [26]. Pressure was increased at ∼0.0069 MPa s^−1^ (1 psi s^−1^) to 0.138 MPa (20 psi) and held until the solution was fully drained, then increased stepwise in 0.069 MPa (10 psi) increments until tunneling—defined as a sustained stream of small (≤ 0.1 mm) bubbles from a single site—was observed, after which pressure was increased continuously at the same rate until cell rupture. All measurements were recorded using a close-range camera (Aven 26700-207 digital handheld microscope).

### D. Statistical analysis

Statistical analyses were conducted using z-tests and one-way analysis of variance (ANOVA) to assess differences in strength, Young’s modulus, and pressure thresholds for tunneling and bursting. When ANOVA indicated significant effects, Tukey-adjusted pairwise comparisons were performed to evaluate differences among groups.

## III. RESULTS

Stress-strain curves of *Chara corallina* cell walls in FIG 1(a) showed high strength in the control samples [63 ± 12 MPa, FIG. 1(b)], a clear reduction in strength after pectin removal (17 ± 5 MPa), and no change after calcium removal or in the shaking controls (59 ± 3 MPa). The control samples had high stiffness, 0.97 ± 0.15 GPa; calcium removal reduced stiffness to 0.63 ± 0.24 GPa, and pectin removal reduced stiffness further to 0.42 ± 0.04 GPa [FIG. 1(c)]. Samples from which pectin had been removed also showed greater stress relaxation under constant strain compared with the control and calcium-removal samples [FIG. 1(d)]. Fracture toughness of the control walls was 9800 ± 2200 J m^−2^ (n=3). We were unable to make repeated measurements after either calcium or pectin removal because the samples were extremely soft and difficult to manipulate. The one value obtained after calcium removal had a toughness value similar to that of the control, while the one value obtained after pectin removal was markedly lower.

Water-permeability measurements showed that calcium removal increased water flux relative to the control, with pectin removal having an even greater effect [FIG. 2(b)]. During gas injection experiments, we identified three distinct modes of gas transport through the cell wall: diffusion, tunneling (a constant and stable stream of large bubbles at fixed locations, Supplemental Video 1 of the Supplemental Material [24]), and sudden efflux through a large crack when the cell ruptured. Tunneling is an advective process that occurs on the time scale of seconds, rather than a diffusive process that forms visible bubbles on the time scale of minutes. The pressure at which gas first tunnels through the wall was 0.98 ± 0.14 MPa in the control samples, and 0.39 ± 0.08 MPa in the pectin-removal samples (p<0.05). As pressure continues to increase, an additional 1-4 nucleation sites were observed. Bursting pressures were consistent with the strengths of the control and pectin-removal groups shown in FIG 3(b), with catastrophic failure occurring at lower pressures (1.1 ± 0.3 MPa) when pectin was removed than controls (1.6 ± 0.2 MPa).

**FIG. 3.**
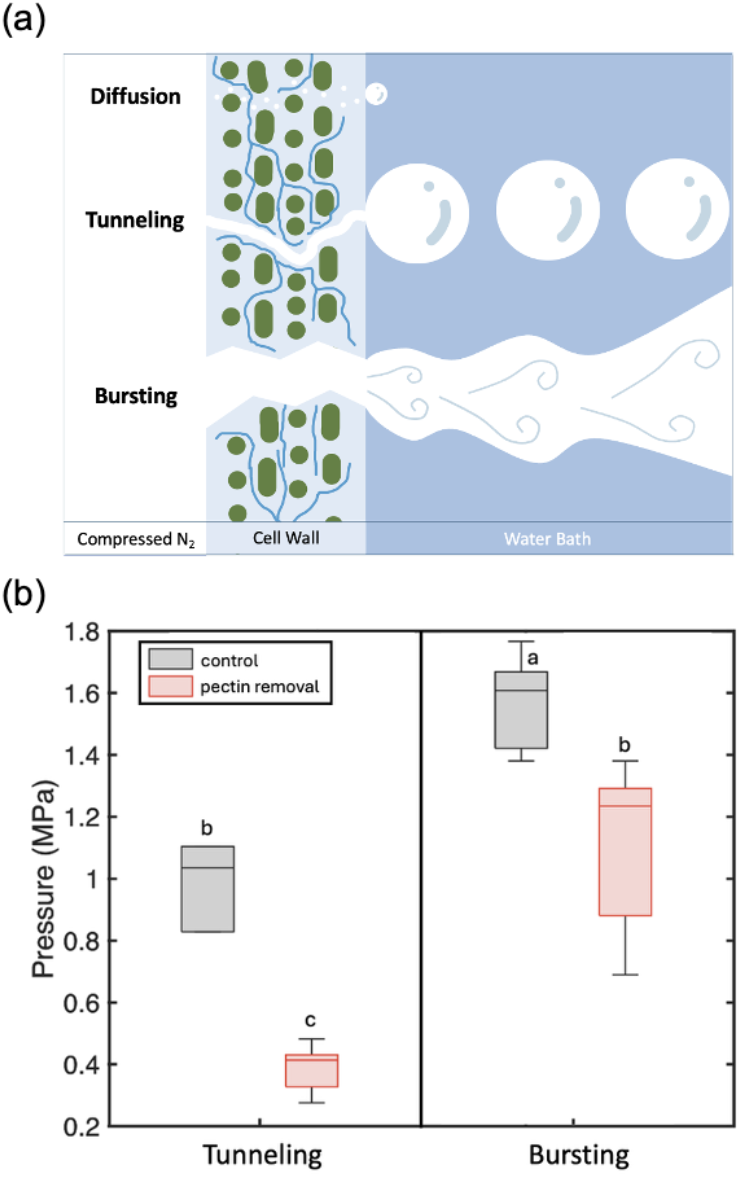
Gas injection experiments. (a) Three modes of gas transport across the *Chara* cell wall (green: cellulose, cross section; blue lines: pectin chains): 1) Diffusion, gas molecules (small white dots) diffuse through the cell walls and accumulate on the outer surface, forming small bubbles; 2) Tunneling, air travels through a pathway that opens under pressure and accumulates to form intermediate sized bubbles that snap off and float away; 3) Bursting, catastrophic network failure resulting in air rushing out of a crack in the cell wall. (b) Pressure threshold for air tunneling through control and pectin removal samples. Lower case letters show statistical differences from Tukey-adjusted pairwise comparisons (one-way ANOVA, p < 0.05).

## IV. DISCUSSION

Pectin is a viscoelastic material with a low shear modulus on the scale of kPa in the frequency range 1– 102 rad/s [13], and until recently was assumed to only affect cell wall porosity [10,12]. However, it is increasingly clear that pectin makes an important mechanical contribution to primary cell walls [9,17-19,20]. Solid-state Nuclear Magnetic Resonance imaging of *Arabidopsis* cell walls shows that pectin polymers are in close proximity to cellulose microfibrils [9], and there are much less signals showing hemicellulose around cellulose. This study challenges the traditional “tethered network” model [8], which assumes that hemicellulose extensively coats and tethers the cellulose network. Mutants with altered homogalacturonan molecular weights show changes in homogalacturonan-cellulose interactions and the relative mobility of both polymers, with lower-MW HG associated with increased mobility and weaker HG–cellulose interactions [18], suggesting that pectin plays a central role in cell wall mechanics via intermolecular interactions. Our study shows that pectin constrains deformation of the cellulose network under both uniaxial and biaxial loading and directly visualizes gas bubble formation driven by gas penetration through a plant-derived material.

### A. Mechanical testing suggests that pectin restricts movements of cellulose under uniaxial loading

Mechanical testing with the Instron system captures the bulk material response of *Chara* cell walls under uniaxial loading. The resulting stress–strain curves exhibit the characteristic behavior of primary cell walls, with an initial stiff elastic regime followed by a more compliant, plastic region. Young’s modulus of the control samples is on the order of gigapascals, consistent with previously reported values [12,22]. In contrast, pectin removal substantially weakens the material (strength) and increases its compliance (reduced slope), in agreement with observations on onion epidermal cells in which pectin digestion resulted in higher incidence of sample breakage during mechanical testing [12]. However, Zhang et al. also report that the Young’s modulus of onion epidermis was insensitive to pectin digestion [12]. The difference between our results and previous observations may arise from the structural organization of onion epidermal tissue, where cells are interconnected by pectin-rich middle lamellae.

The toughness of the control walls (∼10 kJ/m^2^, FIG. S2(a) aligns with values reported by Toole et al for *Chara* [22]. The control samples have higher toughness than typical hydrogels, such as gelatin and alginate, and are comparable to tough double-network hydrogels [27], while lower than insect cuticles [28]. A reduction in toughness after pectin removal, as occurred in our single successful measurement, is consistent with our hypothesis that the cellulose microfibril network becomes easily disrupted when pectin is absent. Stress-relaxation experiments further support this interpretation: samples lacking pectin show greatly increased stress relaxation (the material behaves more like a fluid rather than a solid), indicating that pectin contributes to maintaining microfibril positioning and resisting shear between cellulose elements.

### B. Pectin chemistry affects the local compliance of cell walls under biaxial loading

Pectin is highly branched and has various categories of polysaccharide side chains, including homogalacturonan (HG), xylogalacturonan (XGA), homogalacturonan, rhamnogalacturonan I (RGI), and rhamnogalacturonan II (RGII). HG is a primary pectic polymer consisting of a-1,4-linked galacturonic acids. HG is secreted during cell division and expansion as highly methyl esterified polymers, and is later selectively de-methyl esterified by methylesterases [29]. Demethylated pectate bears calcium-binding moieties. Therefore, chains of HGs can condense by crosslinking with calcium to form junction zones known as "egg-boxes" [30]. The strongest junction occurs between two chains of at least seven unesterified GalA units each.

Our study of *Chara* cell walls shows that removing calcium crosslinks increases water permeability while leaving most bulk mechanical properties under uniaxial loading unchanged, apart from a slight reduction in Young’s modulus. This observation suggests that the effective pore size has increased while increasing the injection pressure for water, and this change occurred locally without affecting the overall mechanical properties.

Consistent with this interpretation, the decrease in Young’s modulus in calcium-depleted samples, despite similar overall strength, indicates increased compliance in the low-stress region of the stress-strain curve (Fig. 1A), likely reflecting greater effective network spacing under biaxial load. A similar pattern was reported by Wang et al. [31], who observed that calcium removal in onion epidermal cell walls had little effect on stiffness but reduced creep (change in displacement under constant load). More broadly, the role of calcium crosslinks in regulating cell wall mechanics is well established in *Chara*: reducing calcium crosslinks promotes wall extensibility under turgor pressure, whereas their reintroduction, along with new pectin deposition, coincides with the cessation of cell expansion [32-34]. Comparable effects have been reported in *Arabidopsis thaliana* [35,36]. Together, these findings, along with our results, support the idea that pectin chemistry, through calcium crosslinking and interactions with cellulose, plays a central role in modulating cell wall compliance, particularly under biaxial loading.

### C. Reduced tunneling threshold due to pectin removal in *Chara* mirrors reduced embolism threshold in vascular plants

Chara cell walls provide a simplified porous structure in which to study the contribution of pectin to the permeability of both water and gas. The increase in water permeability following chemical treatments [FIG. 2(b)] suggests that cellulose and pectin networks are mechanically dynamic and can deform under pressure to facilitate higher volumetric flow, even when a gas-water interface is absent (zero surface tension). The pronounced effect of pectin removal on the tunneling threshold is consistent with the dramatic reduction in fracture toughness, suggesting that cellulose can move more freely without the pectin matrix than in control samples.

Visual observations reveal three distinct modes of gas transport: diffusion, tunneling, and rapid efflux following catastrophic wall rupture. Typical diffusion processes occur in the time frame of minutes, with the rate of diffusion a function of the concentration gradient across the traveled path [37]. The presence of various long polymers imposes steric and frictional hindrance that makes the diffusion pathway longer than in free-form diffusion [38,39]. If the gas molecules interact with the hydrophobic part of the cell wall (cellulose), anomalous diffusion (subdiffusion) would make the transport process slower as well [37]. However, because diffusion takes place within a relatively long timescale, the gas molecules do not distort the cell wall matrix, unlike during gas penetration that occurs on a shorter timescale. During the gas injection experiment reported here, the concentration of nitrogen gas inside of the cylindrical cell wall increases with incrementally applied pressure, and therefore, the rate of diffusion across the cell wall increases as well, leading to an increase in the formation of small gas bubbles on the outer cell wall.

Tunneling produces a steady stream of bubbles that forms within less than a second, clearly distinct from the slower formation of bubbles due to diffusion. During tunneling, the gas finds the weakest point in the cell wall networks. For a 0.1 M SDS solution, the surface tension is ∼39 mN/m, and the corresponding Laplace radius, calculated from the measured average tunneling threshold of controls, is ∼50 nm. This radius is much higher than the typical spacing for the cellulose network (r = 10 nm) based on measurements of atomic force microscopy (AFM) [40,41]. This means that tunneling distorts the cell wall, including both the cellulose and the pectin network. As pressure continues to increase, the appearance of more nucleation sites is consistent with the idea that the global tension in the cell wall has caused more deformation of polymers in the cell wall [FIG. 4(b)-(c)].

**FIG. 4.**
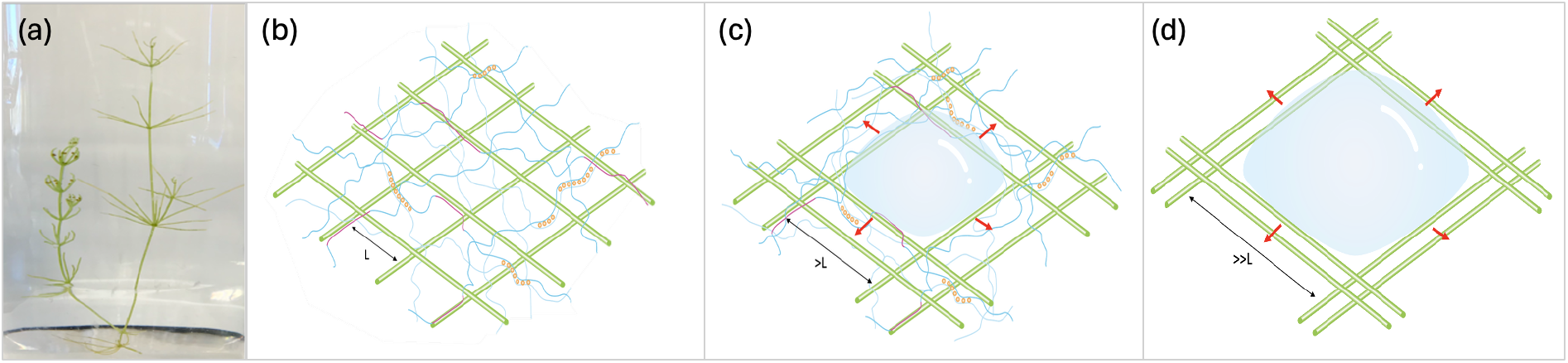
Illustrations of movements of cellulose and pectin networks in Chara cell walls. (a) A *Chara corallina* containing both reproductive (left) and nonreproductive (right) nodes. (b) Schematic representation of the cell wall structure of *Chara carollina* as a cellulose-pectin double network. Cellulose microfibrils are depicted in green, and pectin strands are shown in blue. Orange dots represent calcium ions (Ca^2+^), and the red segments of the pectin indicate regions of close association with cellulose microfibrils. The typical spacing between cellulose microfibrils is labeled as L. (c) Schematic of gas penetration through the cell wall, moving from bottom to top. Gas penetration locally displaces cellulose microfibrils, creating a pore with a diameter larger than the typical network spacing (>L). The pectin network undergoes rearrangement, leading to the breakage of existing calcium cross-links and the formation of new cross-links at different locations. (d) Schematic of gas penetration through the cellulose network without pectin. In the absence of pectin, the cellulose fibers displace easily, and the resulting pore size is much larger than the typical network spacing (>>L).

Meanwhile, based on the plateau regions of the stress-strain curves in FIG. 1(a), irreversible damage in the cell wall networks accumulates as the applied pressure increases. Thus, we hypothesize that in pectin removal experiment, breaking covalent bonds in the pectin network causes cellulose microfibrils to slide easily, thereby creating large holes [FIG. 4(d)]. Since individual fibers of cellulose have strength on the scale of gigapascal [5], the catastrophic bursting of the cell wall likely occurs by pulling cellulose microfibrils out of the pectin network [42].

### D. Biological and material implications

Our findings on gas penetration through *Chara* cell walls are particularly relevant to the functioning of pit membranes, a specialized form of primary cell wall in plant xylem that permits water transport while restricting the spread of air that would otherwise block hydraulic conductivity [43]. During embolism expansion, air invades pit membranes via rapid snap-off events, creating discrete air bubbles. To better understand the material basis that affects embolism expansion in plants, we considered fracture toughness and work of fracture (i.e., the area under the stress-strain curve) of the *Chara* cell wall. Their ratio defines the fractocohesive length, a characteristic length scale that reflects flaw sensitivity [44,45]. When all cracks are smaller than this length, the material behaves effectively as “crack-free.” In the *Chara* cell wall, the fractocohesive length is approximately 0.2 mm, much larger than our estimates of Laplace radii from measurements of gas penetration. We thus interpret the tunneling of gas through the native cell wall as arising from pores locally expanding into stable openings under stress. As an air bubble forms and expands in the cell wall, the surrounding material is toughness enough to stabilize the expansion, rather than tears apart by fracture.

The fractocohesive length of the *Chara* cell wall is also far greater than the typical Laplace radius (∼144 nm) for embolism spread in vascular plants [43]. Because pit membranes are derived from primary cell walls, our results support the idea that embolism spread involves a local modification of the cell wall networks rather than a catastrophic rupture of the entire pit membrane. To date, the process of embolism spread has only been visualized in silica nanochannels under negative pressure, where pore dimensions are substantially larger than those typical of pit membranes [46]. Direct visualization at the level of individual pit membranes remains extremely challenging, in part due to the opaque nature of woody tissues. Our study therefore provides the first direct observation of air penetration through primary cell walls.

It is important to note that pit membranes are more complex than *Chara* cell walls, both in composition and in the spatial organization of their components. For example, lignin—a phenolic polymer—is present in the torus (central region) of pit membranes in some species and provides additional structural reinforcement through covalent bonding within the cell wall matrix [47]. Pectin, in contrast, has been reported in the annulus region of pit membranes, although its presence in the torus remains uncertain [47,48]. During vessel maturation in many species, pectin is partially removed while lignin is deposited [49,50]. Notably, experimental removal of pectin from pit membranes has been shown to significantly reduce embolism resistance [51,53], indicating that pectin plays a key role in limiting air propagation regardless of the precise site of air entry.

Our study highlights an important mechanical principle in plant cell walls: polymer composition and intermolecular interactions can tune mechanical properties of the material long before catastrophic failure occurs [54,55]. Baskin et al. show that anisotropic plant cell expansion depends on mechanical reinforcement of the wall, mainly through cellulose organization, while pectin contributes to the surrounding matrix compliance that enables controlled deformation [56]. This suggests that pectin-cellulose interaction may act as a mechanical regulator, shaping how stresses are transmitted through the wall and how cells maintain structural integrity during growth-related deformation [55,56].

These findings have direct implications for plant mechano-sensing and morphogenesis, where developmental programs are tightly coupled to the perception and interpretation of mechanical signals [55,57]. Plants continuously integrate information on wall strain, turgor-driven tension, and externally imposed forces, and the pectin-rich matrix is increasingly recognized as an active mechanical regulator rather than a passive filler [55,57]. Pectin-mediated control of wall hydration, viscoelasticity, and stiffness gradients may shape how stress cues are distributed across cellulose microfibril networks and subsequently perceived at the cellular scale [54]. Local chemical modification of pectin, such as changes in methyl-esterification, has been shown to directly alter wall rigidity and thereby govern morphogenetic outcomes, including organ initiation and tissue patterning [58,59]. More broadly, this supports the view that pectin remodeling provides a key mechanistic link between nanoscale wall mechanics, mechano-transduction, and emergent developmental form, reinforcing the concept of the cell wall as both a structural material and a signaling platform in plant growth regulation [60,61].

The plant cell wall is a multicomponent composite material made of an interconnected, high-stiffness network and a softer matrix that mediates load transfer and dissipates energy. This combination results in a material with substantial strength and damage tolerance that is difficult to achieve in homogeneous materials [54]. A similar fiber-matrix strategy has inspired the development of synthetic soft materials in which rigid nanofibers or nanocrystals self-assemble within, or co-assemble with, a compliant polymer network, producing hydrogels with enhanced mechanical properties such as increased stiffness and tunable performance tailored to specific applications [62-65]. Furthermore, our study may provide a model for developing water permeable filters that need to withstand high pressure differentials [66].

## ACKNOWLEDGMENTS

The work was supported by National Science Foundation (NSF-MRSEC DMR-2011754) and BSA Graduate Student Research Award from the Botanical Society of America.

## AUTHOR CONTRIBUTIONS

Holbrook, Rockwell, and He conceptualized and designed the research. He conducted most of the experiments and wrote the manuscript. Kim and Chen helped with mechanical measurements in the Suo lab. Svetkey and Rockwell helped perform gas injection experiments.

## DATA AVAILABILITY

The data that support the findings of this study are available from the corresponding author upon reasonable request.

