## Supplemental Figure S1-S4 for "Pectin is a critical contributor to the mechanical properties of *Chara* cell walls"

#### Supplementary Material for
### Pectin is a critical contributor to the mechanical properties of *Chara* cell walls

Zhe He[1*], Junsoo Kim[3], Zheqi Chen[2], Ian D. Svetkey[1], Zhigang Suo[2], Fulton Rockwell[1], N. Michele Holbrook[1]

[1] The Department of Organismic and Evolutionary at Harvard University, MA, USA 02138
[2] John A. Paulson School of Engineering and Applied Sciences, Harvard University, MA, USA 02138
[3] McCormick School of Engineering, Northwestern University, IL, USA 60208

(a) (b)

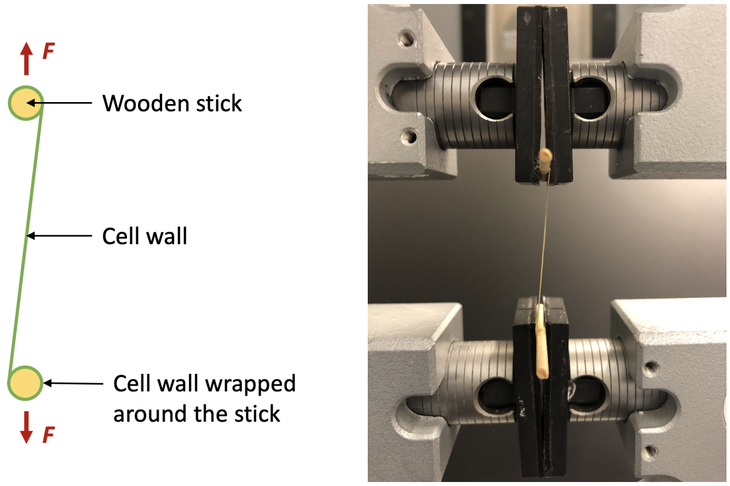


FIG. S1. (a) A schematic diagram showing the cell wall being wrapped around two wooden sticks for tensile tests. (b) A photo mirroring the setup shown in A.

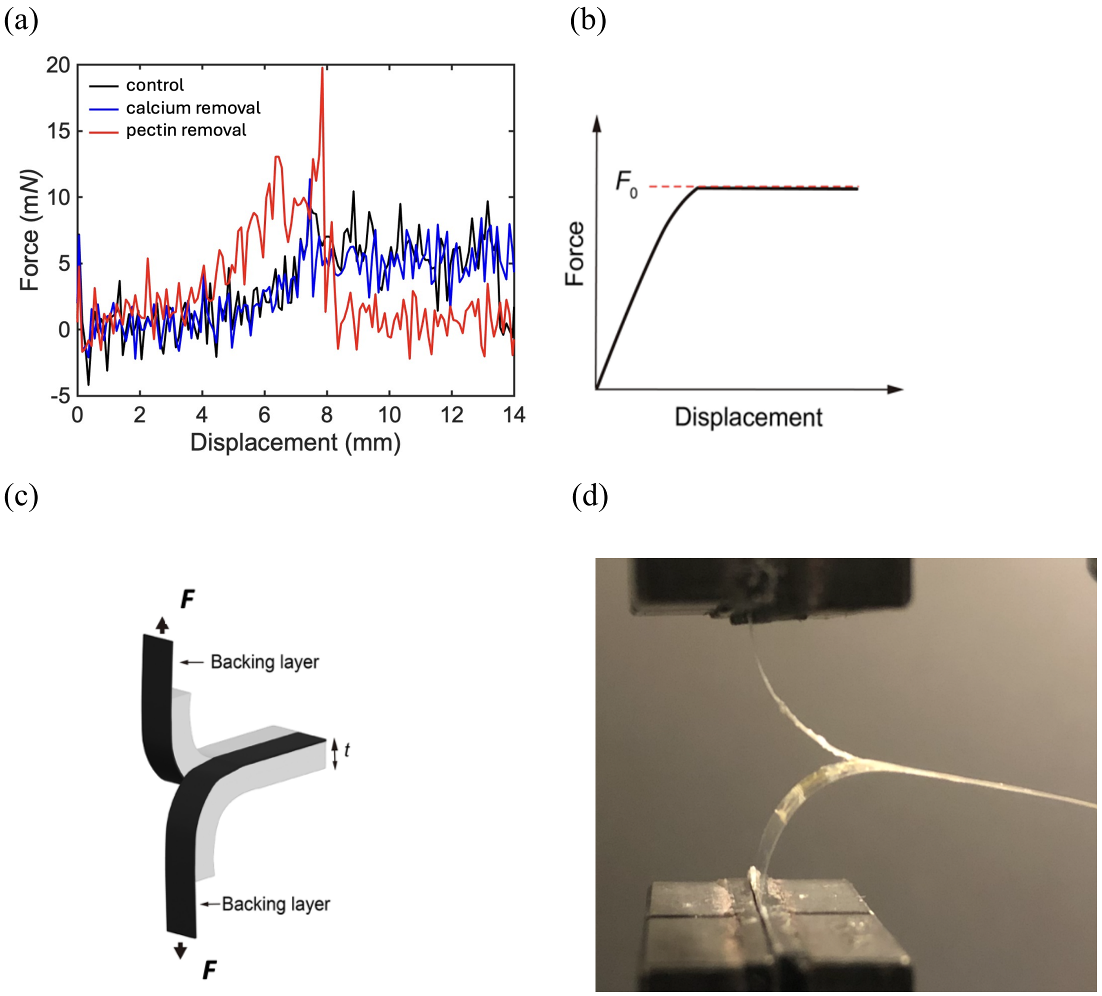


FIG. S2. (a) Force-displacement curves of Chara samples. (b) A schematic diagram showing $F_0$ as the steady-state force for tearing. (c) A schematic diagram of the experimental setup for measuring fracture toughness; (d) A picture of the experimental setup, mirroring C.

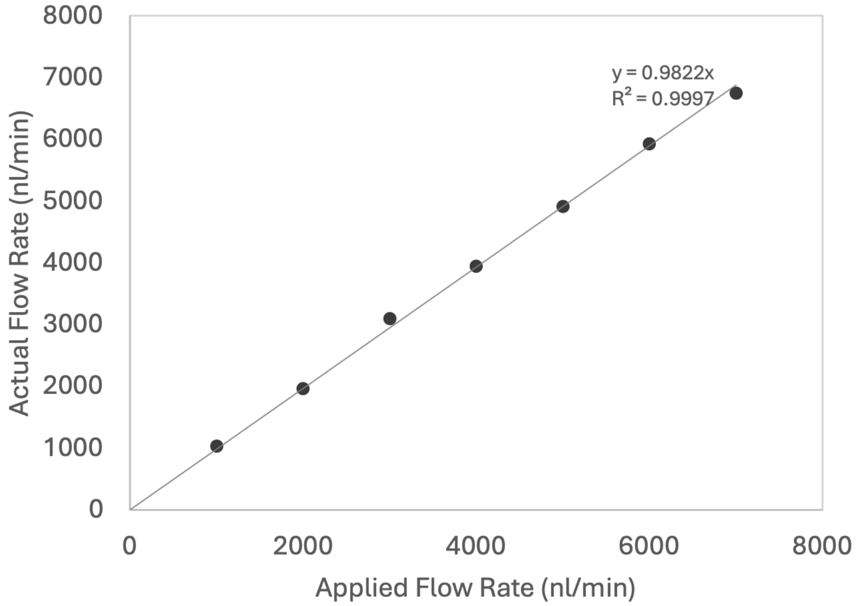


FIG. S3. A calibration curve of the flow meter. The x-axis is the applied flow rate from the syringe pump, and the y-axis is the flow rate measured by the flow meter. linear regression gives a fitting equation of y=0.9822x. $R^2$ = 0.9997.

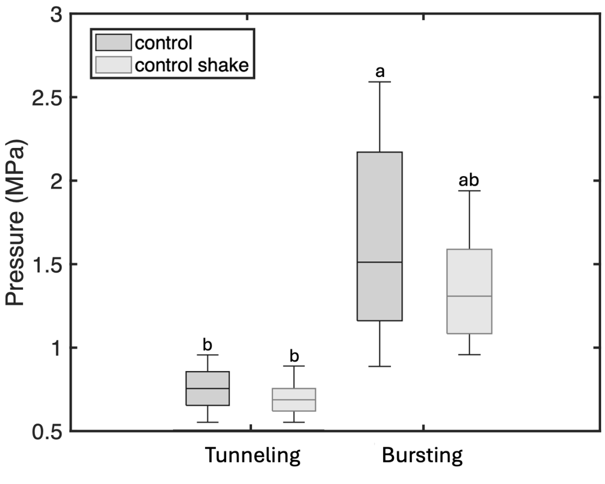


FIG. S4. Pressure threshold for bursting of control and pectin removal samples. Lower case letters show statistical differences, $p < 0.05$.
